# Nutritional Access Modulates Activity of a Small Molecule Enhancer of Endosomal Escape

**DOI:** 10.64898/2026.02.02.703088

**Authors:** Sten Wilhelmson-Andén, Hampus Du Rietz, Hampus Hedlund, Johanna M. Johansson, Wahed Zedan, Anders Wittrup

**Affiliations:** Department of Clinical Sciences, Oncology, Faculty of Medicine, Lund University, Lund, Sweden; Wallenberg Center for Molecular Medicine, Lund, Sweden; Skane University Hospital, Lund, Sweden

**Author notes:** Corresponding Author: Anders Wittrup.

**Keywords:** CAD, nutrient, glucose, siRNA, tumor, chloroquine

## Abstract

For siRNA drugs to be relevant in tumors, poor endosomal escape of these drugs needs to be addressed. Endosomal escape can occur when the endolysosomal membrane is damaged and can be visualized by endogenously expressed fluorescent galectin-9 functioning as damage sensors. Tumor cells have unstable membranes and central parts of tumors have low nutrient levels contributing to reactive oxygen species which can induce membrane damage. We show that nutrient depletion alone does not induce endolysosomal membrane damage in HeLa and MCF7 cells or in HeLa spheroids. Serum depletion, however, enhanced endolysosomal membrane damage in HeLa cells when combined with the membrane destabilizing drug chloroquine, a cationic amphiphilic drug. This effect was almost completely abolished when depleting the cells of glucose, even when serum was present. This phenomenon could not be seen with other cationic amphiphilic drugs like siramesine and loperamide. In a functional experiment, co-treatment with chloroquine and siGFP significantly improved knockdown of GFP in the presence but not in the absence of glucose. Our results have implications for the development of chloroquine as an endosomal escape enhancer of RNA therapeutics in tumor contexts and stresses the importance of considering nutrient levels and tumor size in future screenings.

## Introduction

Today, cancer accounts for almost ten million deaths annually. New therapies, including targeted agents and immunotherapies have complemented surgery, chemotherapy and radiation therapy with improved survival in the last decades. Despite progress, prognosis remains poor for patients with metastatic disease and for several aggressive tumor types, underscoring the need for more effective and personalized treatment strategies.

Small interfering RNAs (siRNAs), i.e., short double-stranded RNAs, that mediate sequence specific knockdown of target RNAs [1, 2], constitute a candidate drug modality for targeted or personalized cancer therapy. siRNAs can silence virtually any gene or oncogene with cytoplasmic mRNA. However, their clinical utility in oncology is currently limited by inefficient cytosolic delivery. After internalization, both ligand-conjugated siRNA molecules and nanoparticle formulated siRNAs are predominantly sequestered in endosomal compartments [3]. For ligand conjugated siRNAs, less than 1% [4] is believed to exit the endosome through spontaneous membrane perturbations – a process known as “endosomal escape”, and enter the cytosol where the RNAi machinery resides and the siRNA can exert its effect. Ligand conjugated and nanoparticle formulated siRNA can achieve efficient hepatic delivery, but reaching extrahepatic tissue, including tumors, remains difficult. Pre-clinical studies have demonstrated efficient accumulation and knockdown in tumors in mice through targeted delivery [5] and through cholesterol-siRNA conjugates [6-8]. However, these strategies require high siRNA doses, reflecting poor endosomal escape. To date, only eight siRNA-based drugs have been approved in the United States, all targeting hepatic indications and none in the field of oncology. Delivering RNA therapeutics to tumors may conferadditional challenges as tumor cells divide rapidly diluting the delivered RNA.Thus, overcoming the bottleneck of endosomal escape and exploiting tumor specific pathophysiological processes will be essential to expand the use of siRNA drugs into the oncology domain [9].

Cationic amphiphilic drugs (CAD) comprise a class of molecules with a weakly basic moiety attached to a hydrophobic region. At neutral pH, the hydrophobic portion permits diffusion across membranes, while at lower pH the basic moiety becomes protonated and the charged molecule can no longer diffuse across membranes leading to accumulation in lysosomes [10]. Once in the lysosomal lumen, CADs neutralize the lysosomal pH and the negative surface charge of intraluminal lysosomal vesicles, leading to the inhibition of luminal hydrolases such as acid sphingomyelinase (ASM) [11, 12]. The resulting accumulation of sphingomyelin (SM) and lysoglycerophospholipids (lysoGPLs) promotes lysosomal membrane permeabilization (LMP) and lysosome-dependent cell death (LDC) [13, 14]. We have previously shown that internalized cholesterol conjugated-siRNA can be released from disrupted endosomal structures by the CADs chloroquine, siramesine, amitriptyline and loperamide, improving knockdown up to ∼50-fold in vitro in HeLa cells [15]. In these studies, we used tumor cells endogenously expressing galectin-9 (gal-9) as a fluorescent membrane damage sensor and visualizing the process as described previously by our group [16].

Although increased lysosomal activity is essential for the elevated metabolic needs of a tumor [17, 18], these changes also make the lysosomal membrane more unstable, exposing a cancer-specific vulnerability that may be exploited therapeutically [19, 20]. Cancers are especially sensitive to LMP [17] and lysosomal cell death pathways [21]. Furthermore, ASM is downregulated in many tumors [22] rendering them more sensitive to CAD mediated membrane destabilization. Central parts of tumors have conditions with low nutrients such as glucose and glutamine, hypoxia and high levels of reactive oxygen species (ROS). ROS is induced by multiple forms of stress including hypoxia (reoxygenation) and nutrient deprivation and is also known to damage lipid membranes through lipid peroxidation [23]. Indeed, one of the most ROS-sensitive organelle is the lysosome due to its lack of common antioxidant enzymes and when the levels of ROS are high, superoxide can cross through and damage the lysosomal membrane [22, 24]. Furthermore, HeLa cells treated with hydrogen peroxide have shown lysosomal leakage of 10 kD fluorescent Dextran (but not 70 kD) after 14 minutes incubation [25].

Based on these observations, we hypothesized that nutrient-deprived tumor cells would exhibit elevated endolysosomal membrane damage, and that nutrient deprivation would synergize with CAD treatment to further destabilize endosomal membranes – thereby enhancing siRNA cytosolic delivery. In this paper we show that nutrient depletion alone is not sufficient to induce endolysosomal membrane damage in HeLa and MCF-7 cells, nor in HeLa spheroids endogenously expressing gal-9. As expected, chloroquine induced endolysosomal damage; however, we observed significantly increased damage when chloroquine treatment was combined with serum deprivation. Strikingly, this synergistic effect was abolished under low-glucose conditions. In functional assays, co-treatment with chloroquine and siGFP enhanced GFP knockdown in glucose-replete conditions but not in glucose-depleted conditions.

Our findings have important implications for the development of chloroquine and related CADs as endosomal escape enhancers of RNA therapeutics in tumor contexts. Our results also highlight the importance of considering nutrient levels and tumor size when designing future screens and evaluating delivery enhancers.

## Materials and methods

### Cell culture and reagents

HeLa and MCF-7 cells were purchased from the American Type Culture collection and were confirmed to be free from mycoplasma. Cells were cultivated in Dulbecco’s modified Eagle’s medium (DMEM, Sigma-Aldrich, St. Louis, MO, USA) supplemented with 10% FBS (Gibco), 100 UmL^−1^ penicillin and 100 mg mL^−1^ streptomycin (Gibco) and 2 mM glutamine (Thermo Fisher Scientific, Waltham, MA USA) at 37 °C and 5% CO_2_. HeLa or MCF-7 cells stably expressing gal-9 fused to yellow fluorescent protein (YFP) (from here on referred to as HeLa gal-9 YFP and MCF-7 gal-9 YFP) were established by transfection followed by antibiotic selection and the establishment of monoclonal cell populations by single-cell seeding as described in previous work [15]. Siramesine fumarate salt (SIR), chloroquine diphosphate salt (CQ), loperamide hydrochloride (LOP) and dimethyl sulfoxide (DMSO) were all from Sigma. Cholesterol-conjugated siRNAs (Accell) were from Dharmacon (Lafayette, CO, USA), with the following target sequences: non-targeting #1: UGGUUUACAUGUCGACUAA; eGFP: GCCACAACGUCUAUAUCAU.

### Medium for nutrient deprivation experiments

Fully supplemented high glucose DMEM medium contained 4500 g/l of glucose, 10% FBS and 1% L-glutamine. For individual nutrient deprivation experiments DMEM medium with 4500 g/l of glucose or with a low glucose content of 1000 g/l was used with or without supplement of FBS and L-glutamine. For combined nutrient deprivation experiments a mix of high-(with FBS and L-glutamine) and low (without FBS and L-glutamine) glucose medium was used in ratios (high:low) of 1:1, 1:3 and 1:9.

### Gal-9 foci quantification

For drug-induced endosomal damage quantification, HeLa gal-9 YFP and MCF-7 gal-9 YFP cells were treated with drugs prepared in DMEM at the indicated times and concentrations. The medium was removed, and cells were washed once with PBS. Cells were fixed with 4% PFA for 10 min at room temperature, washed three times with PBS, and incubated with PBS containing 100 ng mL−1 Hoechst 33342 and imaged with a widefield microscope. CellProfiler was used to detect cell nuclei, cell boundaries, and gal-9 foci in maximum intensity projection images.

### Tumor cell spheroid formation and gal-9 assays

Spheroids were formed by plating 10 × 10^3^ HeLa gal-9 YFP cells (5 x 10^3^, 10 × 10^3^ and 20 × 10^3^ in figure 3) per well in a 96-well spheroid microplate (Corning, Kennebunk, ME, USA) in complete DMEM. For the evaluation of gal-9 foci, the spheroids were allowed to form during 48 hours for nutrient deprivation experiments and in the case of chloroquine treatment for 72 hours plus an additional 24 hours with the drug. For drug treatment, complete DMEM supplemented with 0.1% DMSO or 100 μM chloroquine prepared fresh from stock was added. After 24 h drug treatment, the medium was removed, and spheroids were washed once with PBS before fixation with 4% PFA at 4 °C for 20 min in dark. The PFA was removed, and spheroids were washed three times with PBS. For cryosectioning, spheroids were placed in 15 × 15 × 5mm base molds and OCT Cryomount medium (Histolab, Gothenburg, Sweden) was added and molds were placed at −20 °C for at least 2 h. Approximately 10 μm-thick frozen sections were cut using a Leica CM3050 cryostat, and sections were placed on SuperFrost Plus microscope glass slides (Thermo Fisher). Dako Fluorescence Mounting Medium (Agilent, Santa Clara, CA, USA) supplemented with 5 μg mL^−1^ Hoechst 33342 was applied and a No. 1.5 borosilicate cover glass was placed on top. Spheroids were imaged using a widefield microscope with an oil-immersion objective and x63 lens. Typically, *z*-stacks were acquired with 1-μm intervals, with 10-μm imaging depth for cryosections.

### Drug-enhanced eGFP knockdown

HeLa-d1-eGFP cells [15] were plated in 48-well plates, 3 × 10^4^ cells per well. Unless otherwise stated, incubations with chol-siGFP were performed in OptiMEM for 6 h, followed by drug treatment for 18 h in fully supplemented DMEM with or without glucose. At the end of the experiment, cells were washed with PBS and dissociated by trypsin treatment. Fully supplemented DMEM was added, and the suspensions were transferred to a 96-well V-bottomed microplate and centrifuged at 400 × *g* for 5 min. The supernatant was decanted, and cells were resuspended in PBS, followed by centrifugation again as stated. The supernatant was decanted, and the cells were resuspended in 1% bovine serum albumin (BSA) in PBS for direct analysis using flow cytometry.

### Flow cytometry

For eGFP knockdown analysis, samples were analyzed on an Accuri C6 Flow Cytometer (Becton Dickinson, Franklin Lakes, NJ, USA). The viable population was gated by side scatter/forward scatter evaluation, and the mean fluorescence intensity of duplicate samples was calculated. Mean values of cholesterol-siGFP-treated cells were corrected for background fluorescence by subtracting wild-type cell measurements, and for non-specific treatment effects by normalizing to samples treated with control cholesterol-siRNA but otherwise identical conditions.

### Microscopy

Inverted AxioOberver Z.1 microscope equipped with ×63/1.40 Plan-Apochromat oil-immersion objective lens and Colibri 7 solid state LED light source (Zeiss), MS2000 XY Piezo Z stage (Applied Scientific Instrumentation, Euguene, OR, USA), and an ORCA-Flash4.0 V3 Digital CMOS camera (Hamamatsu Photonics, Hamamatsu City, Japan). The system was used in fast acquisition mode (camera, illumination, and stage) using a SVB 1 signal distribution box (Zeiss) and μCon HS Trigger board (PCIxPress). The system operates under ZEN 2.3 (blue). The following multi-bandpass filters were used (Zeiss): Multiband filter set 92 HE LED (E) with triple-band exciter, emitter, and beamsplitter filters: excitation wavelengths 385, 475, and 590 nm, TBS 405, 493, and 610 nm, TBP 425 ± 30, 524 ± 50, and 688 ± 145 nm. Multiband filter set 90 HE LED (E) with quad-band exciter, emitter and beamsplitter filters: excitation wavelengths 385, 475, 555, and 630 nm, QBS 405, 493, 575, and 653 nm, QBP425 ± 30, 514 ± 30, 592 ± 30, and 709 ± 100 nm. Ten planes were acquired per *z*-stack with 1 µm interval.

### Widefield microscopy image processing and analysis

Images were exported as raw data 16-bit tiff-files from the image acquisition software Zen, and deconvolved using Huygens Professional processing interface and empirical point spread functions. Maximum intensity projections of deconvolved *z*-stacks were created for further analysis in CellProfiler. Spheroids were too large to capture in a single acquisition; hence nine regions were imaged using the tiles function in Zen and put together using the stitching tool. Images were then deconvolved in Zen and maximum intensity projections were created for further analysis in CellProfiler.

### Software

CellProfiler 2.1.1 with customized pipelines was used for gal-9 puncta quantifications, segmentation of cells and nuclei and for layer determination in spheroids (see below). Microsoft Excel for Windows was used for data processing and calculations including normalization of gal-9 foci. In two-dimensional cell culture experiments the individual results were normalized to the negative control (DMSO with full nutrient supplements). In spheroids the individual results were normalized to the whole spheroid. All normalization calculations were performed by dividing by the negative control/whole spheroid. GraphPad Prism 10 for Windows Version 10.3.1 was used to create figures. Final figures were assembled in Adobe Illustrator 2023 version 27.8.1.

### CellProfiler Pipelines

Customized pipelines were made for gal-9 foci quantification in two-dimensional cell culture. Cell boundaries were identified using the YFP signal and nuclei were detected using a watershed function. Nuclei were masked from the cell and gal-9 foci were detected in the remaining cytosolic part of the cell. For spheroids the whole spheroid was detected as an object. Edges in the picture as well as the spheroid edge were cropped 50 pixels to remove light artefacts. The spheroid was shrunk by 200-250 pixels to get a smaller object which in the next step was used as a mask for the whole spheroid. This process was repeated five times, creating an onion-like spheroid with six concentric layers: five outer layers of equal thickness (200–250 pixels, depending on size) and a central layer matched as closely as possible. The area outside of the spheroid was masked (to exclude any debris outside of the spheroid) and gal-9 foci were identified in the six concentric layers.

## Results

### Nutrient depletion does not induce endolysosomal membrane damage

We hypothesized that cancer cells with poor nutritional access are more prone to exhibit endolysosomal damage than cancer cells with sufficient nutritional access. To investigate this, HeLa gal-9 YFP and MCF-7 gal-9 YFP cells were cultured in medium depleted of FBS, glucose and/or glutamine, or as graded combined depletion of the same nutrients to rule out the possibility of a non-linear/U-shaped correlation between nutrient deprivation and membrane damage. In contrast to our expectations, neither nutrient removal nor graded combined depletion led to an increase in membrane damages (Fig. 1a-c).

**Fig. 1:**
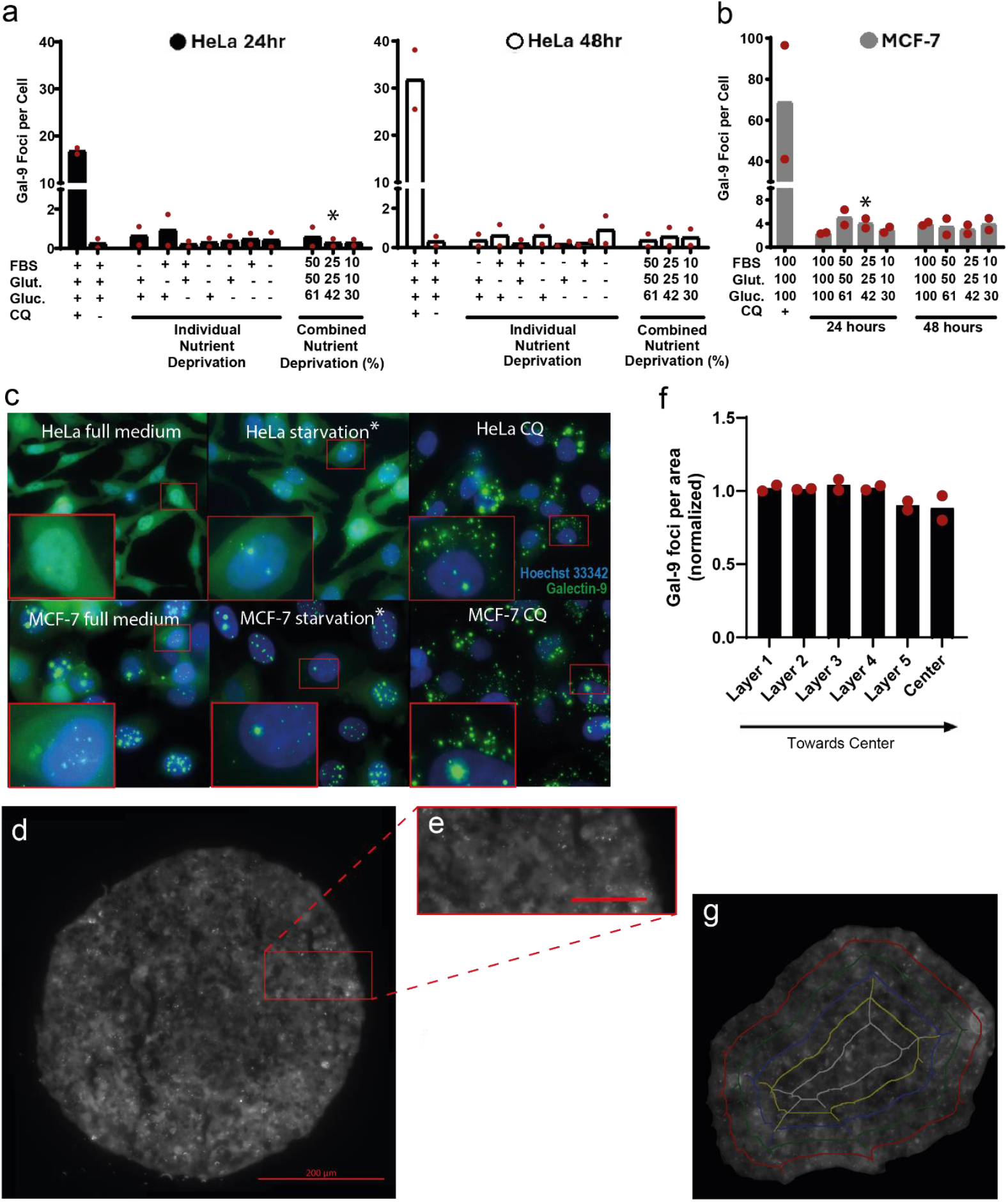
Endo-lysosomal membrane damage is not affected by nutrient depletion. **1a**. HeLa gal-9 YFP cells and **1b** MCF-7 gal-9 YFP cells cultured in medium depleted of fetal bovine serum (FBS), glutamine and/or glucose for 24 or 48 hours (NB discontinued y-axis). Full medium contained 10% FBS, 4500 g/l of glucose and 1% glutamine. As positive control 50 μpM of chloroquine (CQ) was used. The number of gal-9 foci per cell was quantified from widefield microscopy images using maximum intensity projections of a 3 μpm 10-layer z-stack. Individual experi ments were repeated twice. **1c**. Example pictures representative of the conditions full medium after 24 hours, combined nutrient deprivation after 24 hours with 2.5% FBS, 0.25% glutamine and 42% glucose (marked with an asterisk in 1a and 1b) as well as positive control 50 μpM CQ after 24 hours. Note that MCF-7 cells have abundant intra-nuclear gal-9 foci that are unrelated to cytosolic gal-9 foci. **1d**. HeLa gal-9 YFP cells were cultured as spheroids with 10.000 cells per spheroid for 48 hours. **1e**. Enlargement of 1 d, scale bar is 50 pm. **1f**. Gal-9 foci per area was quantified in concentric layers of spheroids and normalized to the whole spheroid (1.0). Red dots indicate two independent experiments with at least four spheroids per experiment. **1g**. Example picture of spheroid divided into concentric layers using Cell Profiler. For details on concentric layer formation and gal-9 foci quantification please see methods section.

Next, we evaluated nutrient-dependent membrane damage in a more physiologically relevant 3D model. HeLa gal-9 YFP cells were cultured as spheroids to create a natural nutrient-deprivation gradient toward the spheroid core. Consistent with our observations in 2D cultures, no increase in membrane damage was detected in nutrient-deprived central regions (Fig. 1d, e), as quantified across concentric layers from the periphery to the spheroid center (Fig. 1f, g).

### The degree of endolysosomal membrane damage induced by chloroquine is dependent on access to nutrients

Although nutrient depletion alone did not trigger membrane damage, we hypothesized that nutrient deprivation may synergize with CAD treatment. To test this, we cultured HeLa gal-9 YFP cells in high- or low-glucose conditions with or without FBS in combination with chloroquine or the vehicle control DMSO. As hypothesized, FBS deprivation in combination with chloroquine treatment did lead to an increase in membrane damage (Fig. 2a, b). However, glucose deprivation did not synergize with chloroquine treatment. In contrast, glucose deprivation almost completely abolished the membrane damaging effect of chloroquine treatment and this inhibitory effect persisted when FBS was also removed.

**Fig. 2:**
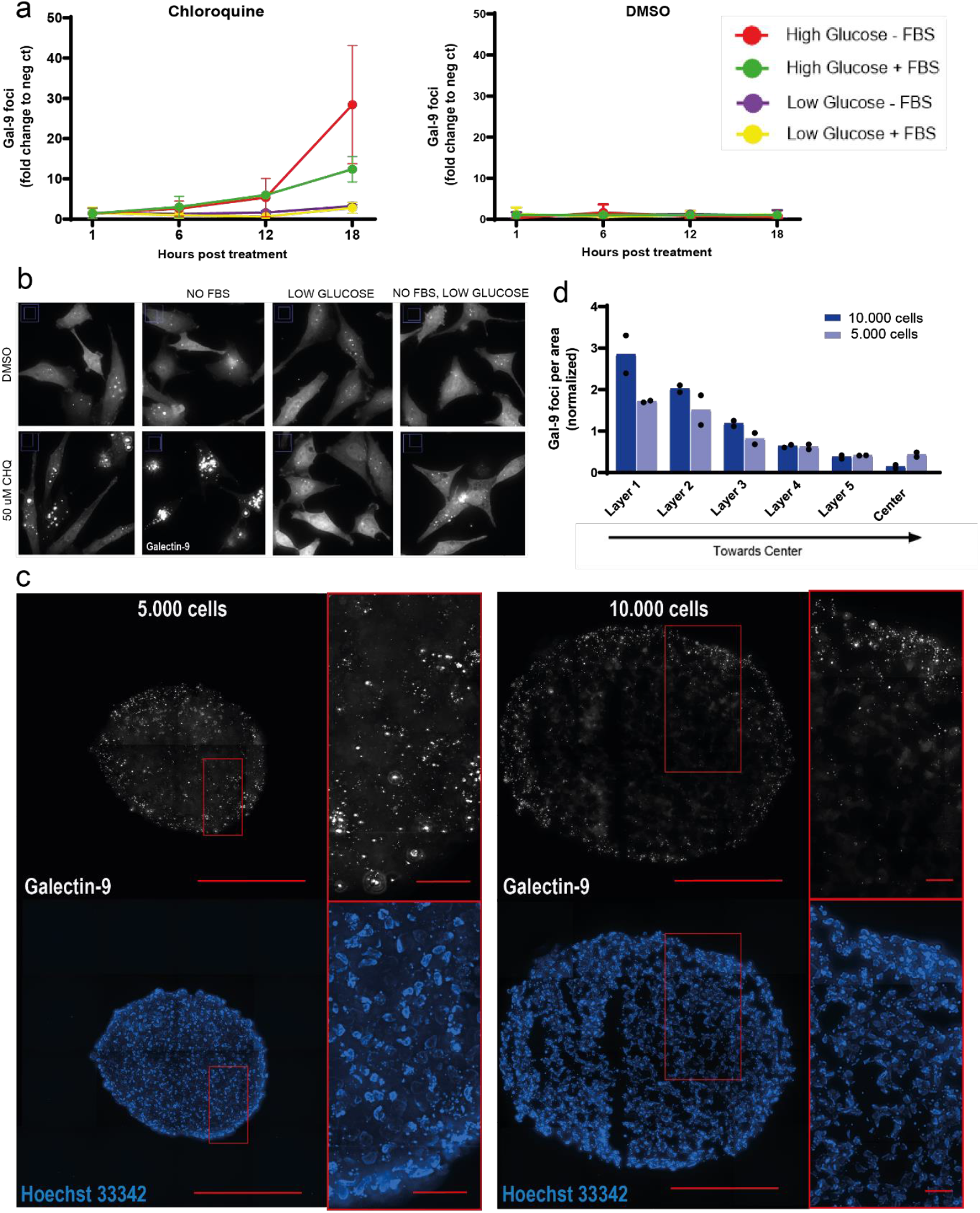
Nutritional access modulates the ability of chloroquine to induce endo-lysosomal membrane damage. **2a**. HeLa gal-9-YFP cells cultured in medium with or without FBS and with low or high glucose and treated with 50 pM chloroquine (CQ - positive control) or DMSO (negative control) for up to 18 hours showing number of gal-9 foci per cell as normalized to full medium (“high glucose + FBS”). Two independent experiments. **2b**. Example pictures from 2a showing widefield fluorescence microscopy images after 18 hours of treatment. **2c**. HeLa gal-9 YFP cells cultured as spheroids with 5.000 (left) or 10.000 (right) cells per spheroid for 72 hours and incubated with 100 μpM chloroquine for another 24 hours and imaged with widefield fluorescence microscopy after cryo-sectioning of the central region and staining with Hoechst 33342. Scale bars are 200 um (50 urn for enlarged areas). Maximum intensity projections of 10-um z-stacks are shown. Images are representative of two independent experiments with three spheroids per condition. **2d**. Concentric layers in HeLa gal-9 YFP spheroids from (2c) quantified with regards to gal-9 foci per area using Cell Profiler (see methods for further details) and normalized to the whole spheroid (1.0). Black dots indicate two independent experiments.

**Fig. 3:**
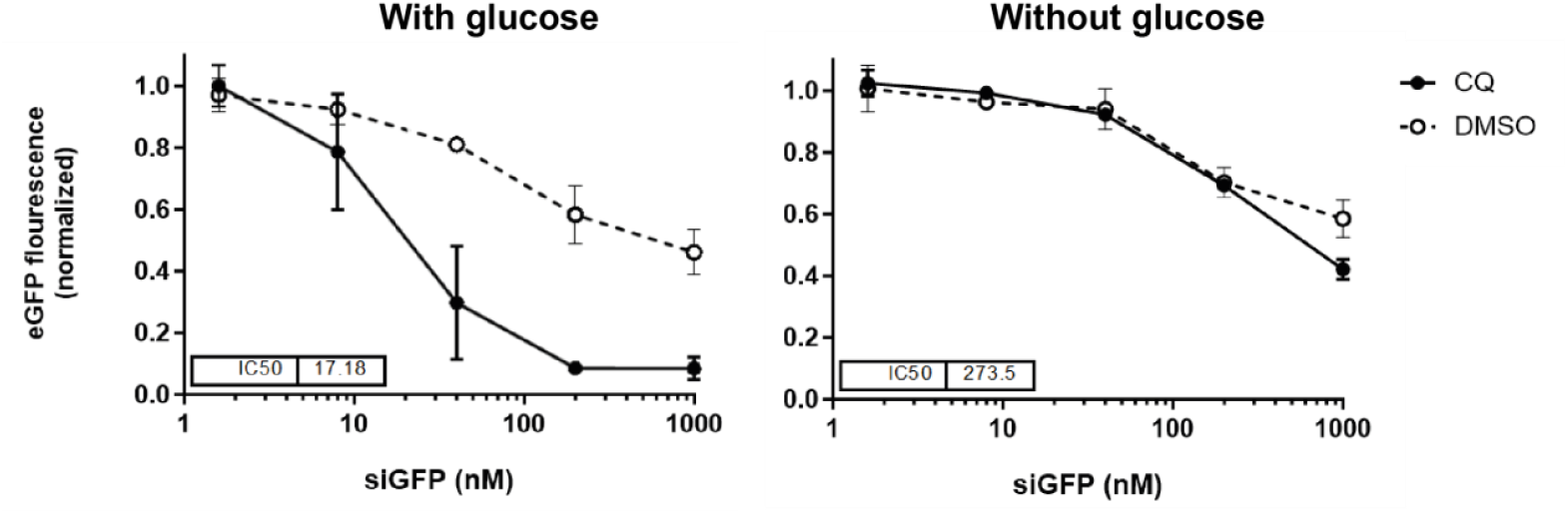
Chloroquine-enhanced chol-siRNA knockdown depends on glucose levels. HeLa-d1-eGFP cells were incubated with the indicated concentrations of chol-siGFP for 6 h, followed by treatment with 60 μpM chloroquine for 18 hours in medium with high glucose content or without glucose. eGFP knockdown was determined by flow cytometry. Circles are mean, and bars are s.d. Data from two independent experiments. Dashed lines indicate DMSO control.

This unexpected finding is consistent with previous work by Gallagher et al. They observed that glucose deprivation abolished the effect of chloroquine on lysosomal swelling in UVW glioma cells, 4TI breast cancer cells and mouse embryonic fibroblasts (MEF) and that blocking glucose flux with 2-Deoxy-D-Glucose significantly reduced chloroquine-dependent lysosome permeabilization, as measured by gal-3 foci formation, both in 4T1 cells and MEF [26].

To verify our results in a more physiologically accurate 3D model, we cultured HeLa gal-9 YFP cells as spheroids to create a natural nutrient-deprivation gradient towards the center. Blood capillaries in tissue typically reside within approximately 100 µm from each other but taking into consideration that tumors are heterogenic in size and vascularity we cultured spheroids of different sizes (Supplementary Fig. S1) and proceeded with 5.000 and 10.000 cells. Spheroids were grown for 72 hours and treated with 100 µM chloroquine for an additional 24 hours. Chloroquine treatment led to a gradient with decreasing numbers of gal-9 foci towards the center (Fig. 2c). The gradient was steeper for larger spheroids than for smaller ones (Fig. 2d), providing further evidence that chloroquine is less effective in deeper tissue. These results are in line with the 2D data and could possibly be explained by low glucose levels in central parts of the spheroids.

### Glucose levels affect the ability of chloroquine to potentiate siRNA knockdown

As low glucose levels inhibit the membrane damaging effect of chloroquine, this could have implications for the potential use of CADs as siRNA delivery enhancers, especially to central parts of tumors where glucose levels are expected to be low. Therefore, we wanted to investigate if the hampered effect of chloroquine in low glucose settings translates into less siRNA knockdown of a target gene. For those experiments HeLa-d1-eGFP cells were treated with chol-siGFP and subsequent chloroquine in the presence or absence of glucose. Indeed, knockdown of GFP was significantly increased by chloroquine in a high glucose setting but this effect was completely abolished in the absence of glucose for low to medium concentrations of siGFP and almost completely abolished for high concentrations of siGFP (Fig. 3).

### Chloroquine analogs do not induce endolysosomal membrane damage

The mechanism as to why the membrane damaging effect of chloroquine is abolished in low glucose settings is unclear. To investigate this further and potentially find chloroquine-like drugs that enhance siRNA treatment even in low glucose/tumor settings, we next explored if analogs to chloroquine could induce membrane damage. The analogs 4-aminoquinoline (AQ) and primaquine (PQ) have been shown to induce de-acidified and swollen lysosomes as well as cell death in glucose deprived conditions [26]. However, treating HeLa-gal-9 YFP cells with AQ and PQ did not lead to an increase in membrane damage regardless of glucose levels (Fig. 4a-c).

**Fig. 4:**
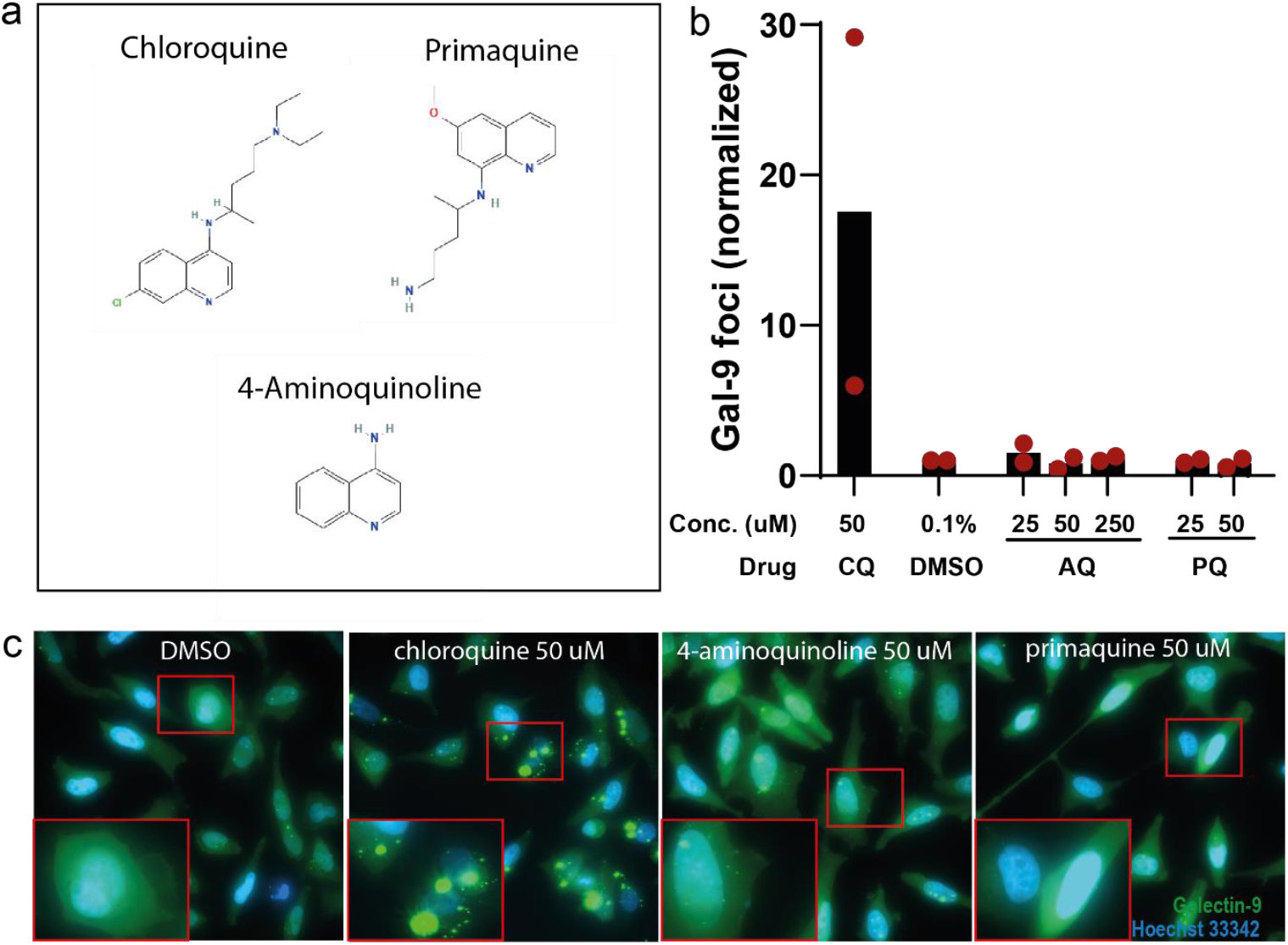
Chloroquine analogues aminoquinoline (AQ) and primaquine (PQ) do not induce endolysosomal membrane damages. **4a**. Chemical structures of chloroquine, 4-aminoquinoline and primaquine from Pubchem at National Institutes of Health. **4b**. HeLa gal-9 YFP cells treated with aminoquinoline (AQ) or primaquine (PQ) in indicated concentrations for 20 hours. The highest concentration of PQ 250 uM led to widespread cell death (not shown). DMSO used as negative control and 50 uM chloroquine (CQ) as positive control. The y-axis shows the number of gal-9 foci per cell normalized to negative control. Red dots indicate two independent experiments. **4c**. Example pictures representative of indicated conditions.

### Other CADs induce endolysosomal membrane damage regardless of glucose levels

Finally, we wanted to explore if the modulating effect of glucose on chloroquine-induced membrane damage is generalizable to other CADs. To investigate this, we evaluated the CADs siramesine and loperamide (Fig. 5a), which have previously been shown to induce membrane damage conducive to siRNA delivery [15]. As expected, treating HeLa gal-9 YFP cells with siramesine or loperamide did induce membrane damage, but the extent of damage was the same regardless of glucose levels (Fig. 5b,c). Thus, the modulating effect on chloroquine-induced membrane damage by glucose availability does not appear to be generalizable to the entire molecular class of CADs, as evidenced by the CADs siramesine and loperamide being insensitive to glucose depletion.

**Fig. 5:**
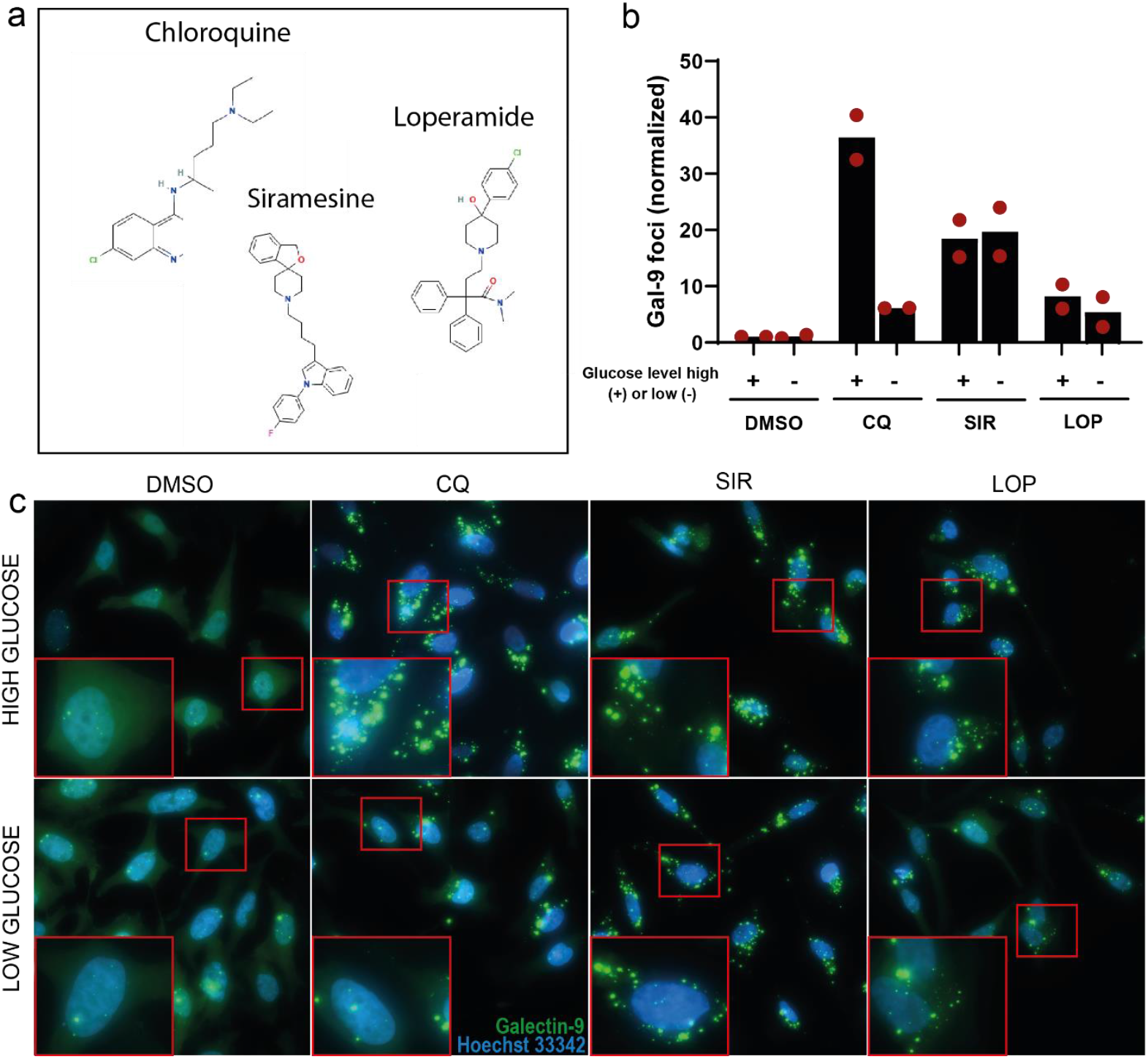
Siramesine and loperamide induce endolysosomal membrane damages indiscriminately of glucose levels. **5a**. HeLa gal-9 YFP cells treated with 10 μpM siramesine (SIR) or 20 pM loperamide (LOP) for 20 hours. DMSO was used as negative control and 50 μpM chloroquine (CQ) as positive control. Y-axis shows gal-9 foci per cell normalized to negative control with high glucose levels. Red dots indicate two independent experiments. **5b**. Example pictures representative of indicated conditions. **5c**. Chemical structures of chloroquine, 4-aminoquinoline and primaquine from Pubchem at National Institutes of Health.

## Discussion

We show that nutrient depletion does not lead to an increase in endolysosomal membrane damages as visualized by gal-9 foci in HeLa cells, HeLa spheroids or MCF-7 cells. One explanation is that cancer cells have become highly adapted to nutrient deprivation and that vesicle membrane integrity therefore is not compromised by deprivation. Another possibility is that small damages do occur but are repaired before they are large enough to expose glycoproteins to the luminal side of vesicles and attract galectins. Recruitment of the endosomal sorting complex required for transport (ESCRT) repair system precede the recruitment of galectins and interference with this system makes otherwise reversible lysosome damage lethal [27]. However, such small and rapidly repaired disruptions are unlikely to permit siRNA leakage. Indeed, for LNP-delivered siRNA, our group has previously shown that galectin-positive damage can be indicative of functional cytosolic release [28].

Although nutrient depletion alone did not trigger membrane damage, we found that serum depletion synergized with chloroquine in inducing membrane damage. In cancer cells, serum deprivation leads to an increase in ROS and oxidative stress [29, 30] which in turn can damage lipid membranes through lipid peroxidation [23]. However, tumor nutritional deprivation affects numerous pathways [31] including apoptosis [32-34] and autophagy [33]. Autophagy protects cells when deprived of nutrients and cells die by apoptosis when autophagy is pharmacologically inhibited [35, 36]. Indeed, all clinically relevant CADs, including chloroquine [37], neutralize the lysosomal pH effectively and thereby inhibit autophagic degradation. Consequently, we do not know if the increased amount of membrane damage seen in concomitant serum deprivation and chloroquine treatment is an additive effect that can be exploited therapeutically or if it merely represents a pre-apoptotic cell stage.

Further investigation is needed to evaluate its significance in the context of improving siRNA delivery.

While serum depletion potentiated the effect of chloroquine in inducing membrane damage, glucose deprivation abolished the membrane damaging effect. The fact that this inhibitory effect persisted when depleting the cells of FBS, together with the results from spheroids where diminishing trend of membrane damage towards central parts of chloroquine treated spheroids were seen, indicates that the abolishing effect of glucose deprivation is more important than the effect of serum. However, the ability of chloroquine to penetrate the spheroid can also contribute to central regions exhibiting less membrane damage. Another possible contributing factor could be that extracellular pH, which is lower in central parts of spheroids, result in more protonated chloroquine extracellularly and hence less intracellular uptake. Nonetheless, the neutering effect of glucose is clear and corroborated by data from Gallagher et al. [26]. Though the underlaying mechanisms remain to be elucidated, the phenomenon suggests that chloroquine might be a sub-optimal choice as an inducer of membrane damage and siRNA delivery enhancer to tumors where glucose levels are known to be low, at least in central parts. This is supported by previous work from our group where we showed greater knock down in GFP signal in outer layers of siGFP/chloroquine treated spheroids [15]. This may have implications for the development of endosomal escape enhancers, where chloroquine has been one of the most studied compounds [38].

While chloroquine was shown to be glucose dependent, siramesine and loperamide are effective in inducing membrane damage (though less so than chloroquine in high glucose settings) in HeLa cells also in conditions with low glucose levels. However, previous work from our group has shown that siramesine and loperamide, despite inducing widespread membrane damage, are unable to potentiate siRNA knock down in 3D tumor spheroids.

Additionally, the membrane damaging effect of loperamide is highly cell type dependent [15]. Thus, their failure as delivery enhancers cannot be attributed to glucose levels, and other mechanisms limit their efficacy.

Given that cancer cells often exhibit altered lysosomal morphology and vulnerabilities, synergistic strategies may offer a path forward. Combinations of agents that perturb endo-lysosomal membranes through distinct mechanisms, such as CADs with ursolic acid [39] or ferroptosis inducers like salinomycin or disulfiram that promote lysosomal iron accumulation and ROS generation [40, 41], could enable a cancer cell targeted strategy to enhance RNA delivery. Additionally, certain cytostatic drugs including vincristine increase the size of lysosomes and induce apoptosis-like cell death preceded by LMP in HeLa cells. Synergistic effects were seen when treating HeLa cells with both vincristine and siramesine [21]. Hence, the challenge will be to find combinations with a sufficient therapeutic window where RNA delivery is enhanced but cytotoxicity, at least to healthy cells, are minimized.

## Acknowledgements

No acknowledgements to be made.

## Author Contributions

A.W. conceived the study and supported funding. S.W., and A.W. designed the experiments. S.W. performed the experiments with support from H.H and J.M.J. S.W. analyzed the data and created figures. H.D.R. provided support and supervision throughout the study. S.W. and A.W. wrote the manuscript with input from all authors.

## Statements and Declarations

### Ethical considerations

Experiments conducted for this article comply with current Swedish law. No ethical approval was required.

### Consent for publication

Not applicable.

### Data Availability

All data are present in the paper and/or the Supplementary Materials. Raw data related to this paper may be requested from the corresponding author.

## Supplementary

**Figure S1.**
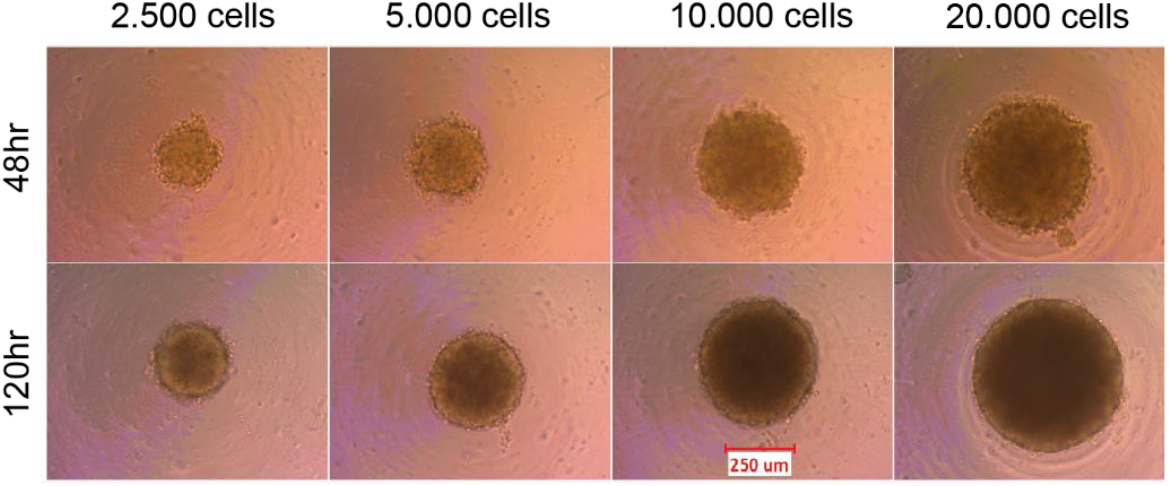
HeLa spheroids of varying sizes. HeLa gal-9 YFP cells cultured in full DMEM medium in round bottom spheroid plates for 48 or 120 hours with indicated cell numbers and visualized with light microscopy. Scale bar 250 um.

