## Supplementary Fig. S1 for "Nutritional Access Modulates Activity of a Small Molecule Enhancer of Endosomal Escape"

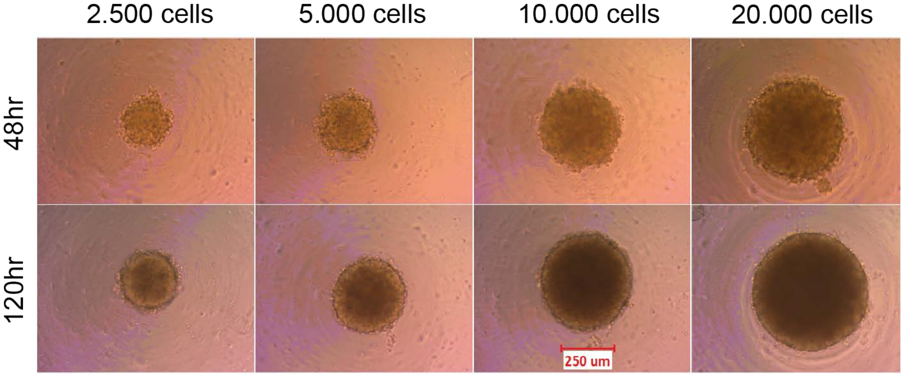


**Figure S1. HeLa spheroids of varying sizes.**
HeLa gal-9 YFP cells cultured in full DMEM medium in round bottom spheroid plates for 48 or 120 hours with indicated cell numbers and visualized with light microscopy. Scale bar 250 um.
